# A framework to assess the contribution of bioaerosols to the outcome of meteorological contexts favorable for rainfall

**DOI:** 10.1101/070532

**Authors:** Cindy E. Morris, Samuel Soubeyrand, E. Keith Bigg, Jessie M. Creamean, David C. Sands

## Abstract

Rainfall feedback results from the sensitivity of atmospheric processes to environmental conditions that are generated by a preceding rainfall event. Feedback that is persistent over several weeks is most likely due to environmental phenomena that involve growth and therefore most probably involves aerosols of biological origin. Based on a tool developed to quantify feedback at specific sites from historical daily rainfall data and maps of the feedback trends (http://w3.avignon.inra.fr/rainfallfeedback/index.html) we have generated a series of site-specific and season-specific hypotheses about the extent to which aerosols – from biological sources in particular - influence the outcome of meteorological conditions that are favorable for rainfall. We illustrate how the tools we report here and elsewhere can be applied in a framework of rationale for the design of field experiments finely tuned to site-specific hypotheses and thereby to a more refined understanding of the contexts of geography, season and land use that underlie the extent to which aerosols influence the fate of cloud processes.

---

Rainfall is critical for water availability. Anticipating rainfall is vital for flood forecasting and management of urban drainage (39, 40); for filling catchments (24) essential for aquatic wildlife, drinking and irrigation water, and electricity generation; and for the planning of planting of crops. Changes in rainfall patterns have numerous consequences including but not limited to altered primary production and the timing of leaf and fruit development (42), declines in the viability of herds of domestic animals (2), socio-economic phenomena that can culminate in civil conflict (8, 11), and contribute, in extreme cases, to the collapse of civilizations (38). Management, adaptation to, and mitigation of the effects of drought for example, as in the current dire situation in California (1), depend on accurate forecasts of precipitation and knowledge of how human activities influence precipitation.

Rainfall patterns depend on synoptic-scale atmospheric circulations. However, the atmosphere is never free of the aerosols that can influence the outcome of meteorological phenomena. Therefore, it is easy to understand that their decisiveness in the processes leading to rainfall is still under debate. There is need for new approaches to address the question of when and where aerosols can make or break the outcome of meteorological conditions that are ripe for rainfall. This need comes hand in hand with the increasing awareness that human activities that generate aerosols (21) and changes in land cover (32) have a marked impact on precipitation because they influence the conditions under which aerosols operate (e.g., temperature and the availability of moisture) and the types and abundance of aerosols produced.

How can the extent to which aerosols influence precipitation be disentangled from the effects of the prevailing meteorological conditions if these two factors are always associated? We recently developed a tool that quantifies rainfall feedback, the apparent sensitivity of rainfall to aerosols that are generated and released into the atmosphere after rainfall events of a certain intensity (28). This tool was developed from observations of feedback patterns that persisted for several weeks in daily rainfall data and similar patterns of increased concentration of ice nucleation active aerosols after rainfall (3, 5). It is difficult to explain such persistent phenomena in terms of physical processes such as increased soil moisture, splashing of aerosols from impaction of raindrops, or trends in ambient meteorological conditions. In light of the numerous particles of biological origin (bacteria, fungi, etc.) in the atmosphere that are active as cloud condensation nuclei (CCN), giant CCN (GCCN) and ice nuclei (INP), their emissions from vegetation and their persistent increase after rainfall (described in detail in the paper that describes the tool to calculate rainfall feedback (28)), we feel that hypotheses about the roles of these biological aerosols in precipitation need to be elaborated and assessed.

With the tool we developed to assess the intensity of rainfall feedback (28) we produced maps of regional and seasonal trends of rainfall feedback in western USA (http://w3.avignon.inra.fr/rainfallfeedback/index.html). The many factors that could influence rainfall feedback are likely to vary among geographic sites in terms of how the atmospheric conditions at these locations promote the increase of cloud-active particles, their emission, aging and interaction with other aerosols. By characterizing the relationship of land use, topography and seasons to rainfall feedback, we have created a set of hypotheses about the extent to which aerosols influence precipitation at specific locations and during different seasons. It should be noted that these are hypotheses, i.e. they are the fruit of speculation and the foundation for future empirical studies. By creating and sharing hypotheses, the pertinence of experimentation can be optimized and results can be validated across comparable sites or conditions.

## MATERIALS AND METHODS

Trends in rainfall feedback at the 1250 sites in the western USA available for this study were identified as described previously (28). We paid particular attention to the location of regions with the greatest intensity of positive and negative values of the feedback index (***F***) and of marked differences between the spring-summer (April-Sept) and fall-winter (Oct.-Mar) seasons. Hypotheses were generated based on topography, knowledge of prevailing atmospheric circulation and land use history of the regions.

The hypotheses were also generated based on knowledge of sources of aerosols and their dynamics. Briefly, cloud droplets can grow to form raindrops by i) coalescence aided by CCN and high concentrations of GCCN or ii) the Bergeron process that is catalyzed by INPs at temperatures below 0°C depending on the temperature at which the INPs are capable at catalyzing this process. In addition to the regularly occurring emission of these three types of aerosols from various natural and anthropogenic sources, surges of emission of GCCN and IN of microbiological occur after rainfalls (19). The nature of the ensemble of aerosols, their abundance in clouds and their interactions can lead to various outcomes for precipitation. For example, microbial IN catalyze ice formation at temperatures much warmer than nearly all other naturally-occurring IN (25) and in particular between -3°C and -8°C where the Hallett-Mossop (H-M) process leads to an ice crystal multiplication. Hence, surges of emission of microbial IN may greatly enhance the probability of precipitation, or – if too many crystals are produced - may inhibit it due to competition among ice crystals for the available water. The H-M process requires the presence of cloud drops >24 µm in diameter that could be assured by the associated surge in emission of microbial GCCN (16). In stratiform clouds, high concentrations of GCCN can lower water content and the possibility of rain by producing drizzle that does not reach the ground. An over-abundance of CCN can be inhibitory to precipitation (35). Under specific conditions, rainfall is followed by rapid increases in new particle formation in the atmosphere (4, 6) that have been shown to act as CCN. Overall, the combination of the regularly-occurring aerosols with rain-responsive aerosols can have outcomes that can lead to either more favorable or inhibitory conditions for subsequent rainfall thereby setting the stage for feedback.

## HYPOTHESES

### Hypothesis 1. Orographic precipitation is propitious for positive rainfall feedback

A map of the sites with the greatest positive values of ***F*** (> 0.5) (see Fig. 4 of Morris et al (28)) suggested that orographic precipitation was a predisposing factor for positive feedback. The position of these sites was in marked contrast to the location of the sites with near-zero values of ***F*** that were mostly in the plains east of the Rocky Mountains.

The short residence time of an air parcel within an orographic cloud means that precipitation is much more dependent on the efficiency or speed of development of precipitation than in non-orographic clouds (20). This suggests that orographic precipitation would be particularly sensitive to changes in concentrations of efficient INPs and GCCN more so than precipitation from convective uplift of water vapor and aerosols. Such phenomena have been corroborated by modeling (20). Additional corroboration of the influence of efficient INPs on orographic precipitation comes from field observations showing that INPs active at temperatures warmer than -10° C are lost early in the precipitation history of orographic clouds (37). Furthermore, these observations suggest that the vegetation upwind of the mountain ranges mentioned above would be important sources of rain-responsive cloud-active aerosols.

It should be noted that precipitation in Washington, Oregon, California, and Arizona is largely impacted by atmospheric rivers, which are concentrated and narrow meridional bands of water vapor that are transported from the tropics and make landfall along the west coast of the USA (7, 29, 30). These are the general overall trends for these regions and specifically on the mesoscale and for systems generated over/fueled by Pacific circulation patterns. However, it should be kept in mind that the North American monsoonal circulation and storms generated from easterly flow along the Colorado Front Range can sometimes offset this trend. Nevertheless, the Pacific Northwest coast (Washington and Oregon), like California, is also a site where orographic clouds form. However, ***F*** is strikingly more negative in the Pacific Northwest than in California. There might be important differences in the types and seasonality of aerosols in these regions due to vegetation, as discussed in more detail below, and also due to the numerous pulp and paper mills, large sawmills with wood waste burners, and aluminum smelters in the Pacific Northwest. The paper mills in particular produce prolific amount of GCCN (up to 10^19^ / sec) that enhance precipitation downwind (18) thereby possibly masking or confounding effects from rain-responsive aerosols. The hypotheses concerning orographic precipitation and feedback could be addressed by conducting focused field studies in these orographic cloud formation regions to disentangle the effects from orography and aerosols. An interesting approach would be to compare similar measurements of meteorology, aerosols, and precipitation in disparate mountain ranges, which include regions with variable aerosol sources.

### Hypothesis 2. The impact of vegetation on rainfall depends on its biodiversity, phenology and health

Another trend that we reported was the marked negative feedback across much of the Pacific Northwest. Furthermore, when we examined the seasonal effect of feedback, there was a distinct transect from the Pacific Northwest to the southeastern part of the USA. The Pacific Northwest was characterized by high positive feedback in the spring-summer season and negative in the fall-winter season whereas the southeastern part of the USA showed the inverse pattern. There was a transition of the trend along the western edge of the Rocky Mountain Range (Fig. 5, Morris et al (28)).

This raises the question about the seasonal changes at these sites that would be favorable to rainfall during one season and inhibitory to rainfall in another season. An interesting hypothesis arises from the observation that nearly half of the sites with high positive rainfall feedback in the fall-winter season and negative feedback in the spring-summer are in the wheat belt of the Great Plains. In the USA wheat belt states, more than 30 million ha are planted to wheat (of which half is winter wheat, planted from September to October) and are harvested from May to July. Since the late 1800's wheat in the USA has been subject to regular epidemics of stem rust (caused by the fungus *Puccinia triticina*) and leaf rust (caused by *P. graminis* f. sp. *tritici*) (9, 17). The peak of the epidemics generally occurs in summer months with up to 10^13^ urediospores of the fungus emitted into the air per 5 ha of field per day or per hour of combining at harvest (12, 31). As mentioned above, the air-borne spores of these and other rust fungi (urediospores) are efficient INPs active at temperatures as warm as -4° C and it has been proposed that such prolific emissions could overcharge clouds with INPs thereby having inhibitory effects on the formation of raindrops (27). Under this scenario, rainfall in the fall-winter season would enhance growth of cold-tolerant ice nucleation active microorganisms such as *P. syringae* on winter wheat and other vegetation that could have favorable impacts on subsequent rainfall because background concentrations of such aerosols would be relatively low. This would be in contrast to the spring-summer effect of prolific urediospore production that would be inhibitory to rainfall formation.

The nature of the vegetated land cover in the Great Plains is markedly different from that of the Pacific Northwest where seasonal rainfall feedback trends are also the inverse of the trends in the Great Plains. This leads us to speculate that the properties of the Pacific Northwest vegetation in winter foster production of cloud-active aerosols that inhibit cloud drop growth and raindrop formation in a scenario analogous to what we proposed for the Great Plains in the summer. Approximately 25% of the land cover in the Pacific Northwest is composed of evergreen coniferous forests and an additional 25% is composed of mixed conifer and deciduous hardwood forests (15). Only about 45% of the land use in this region constitutes crops, pastures, grasslands and steppe. Therefore, during the winter months, foliage of conifers would be dominant over that of annual and deciduous plants. Although there are important epidemics of rust diseases on conifers in the Pacific Northwest that could be sources of INPs, production of urediospores in the disease cycle does not occur in the fall-winter season (13). Furthermore, the bacterial flora on leaves of conifers and other gymnosperms has an unusual genetic structure, dominated by species in the Bacteroidetes phylum and only rarely harboring ice nucleation active bacteria whereas those on angiosperms are dominated by Proteobacteria and frequently harbor ice nucleation active bacteria (23, 33).

The seasonal dichotomy of rainfall feedback in the Pacific Northwest might result from environmental chemical conditions conducive to NPF. The abundance of these new particles can vary across seasons, and in some forested sites the peak abundances can occur between November and March (34). Rapid increases in NPF also typically occur after rainfall because rain washes out other particles that compete for the condensation of the gases (6). But, as mentioned above, persistent increases of new particles likely due to rain-induced growth of soil microorganisms have been observed in forests (4). In light of these observations we suggest that the wet and mild winter conditions in the Pacific Northwest forests are conducive to production of quantities of cloud-active aerosols that, when added to the pool of existing aerosols, are abundant enough to serve as CCN and potentially inhibit precipitation. Overall the effects of land use on rainfall feedback evoke an ensemble of hypotheses concerning the dynamics of biological and biogenic cloud-active aerosols. Field studies to address these hypotheses could attempt to characterize more specifically the biological aerosols – in terms of the identity of the microbial species - and the biogenic aerosols involved and the land cover from which they originate. Not only will it be useful to elucidate the dynamics of abundance of the biological aerosols to test the hypotheses mentioned here, but it will be important to identify the periods when upward flux of such particles could occur. For example, rain-responsive ice nucleation active bacteria such as *P. syringae* tend to become airborne in warm sunny conditions with wind strengths exceeding 1 m/s (22), conditions that are not usually associated with rain. There is much that is to be learned about microbial flux given the important paucity of direct measurements (26).

### Hypothesis 3. Urban centers favor positive feedback

In a previous study on rainfall feedback in Australia (5), sites in large urban centers and downwind had positive rainfall feedback. Here, the values of ***F*** in the immediate vicinity of San Francisco, Los Angeles and San Diego, California; Phoenix, Arizona; Salt Lake City, Utah and Denver, Colorado tended to be greater than those of the more distant surrounding area. Some of these sites might be influenced by orographic processes as described above, but the apparent downwind trends suggest a role for additional factors such as urban aerosols, which have mostly been considered as inhibitory to rainfall (35). We were able to evaluate the downwind footprint of the San Francisco and Phoenix urban centers because there were sufficient downwind sites in contrast to the settings of the other large urban centers that are not importantly impacted by orographic precipitation. Furthermore, the San Francisco and Phoenix regions are unencumbered by large point sources of anthropogenic pollutants such as ozone, volatile organic compounds (VOCs), NOx, and SO_2_, which can create secondary aerosols that serve as CCN and delay the onset of precipitation (14, 36). Both of these urban centers had downwind plumes of positive ***F*** for distances of 200 to 300 km. The contours of the plumes did not mimic the topography and were consistent with an urban plume extending in the downwind direction (Fig. 1). The overall pattern of ***F*** values around Phoenix is in marked contrast to that in the adjacent region of Tucson which is also on the windward edge of the Mogollon Rim of the southern Rocky Mountains and which are both primarily influenced by incoming air masses from the southwest^1^. Phoenix and Tucson are markedly different in terms of both their physical and population sizes as urban centers and in terms of the surrounding and downwind land use. The regions around Phoenix and Tucson also had different seasonal dichotomies in rainfall feedback with the inter-seasonal variability for the Phoenix region being greater and more markedly significant (mean ***F**_Oct-Mar_* = 0.53, ***F**_Apr-Sept_* = -0.43, pairwise t test p = 0.0001) than for Tucson (mean ***F**_Oct-Mar_* = 0.19, ***F**_Apr-Sept_* = -0.19, pairwise t test p = 0.0367) (based on 8 to 9 sites up to 115 km north of each of the cities and eastward to the edge of the Mogollon Rim or up to 200 km eastward in the case of Tucson). Urban centers are known to emit large quantities of particulate matter depending on the size of the urban center (36) and some of these have negative effects on rainfall (35). However, the trends in rainfall feedback suggest that aerosols from urban centers are somehow favorable to rainfall feedback.

**Figure 1.**
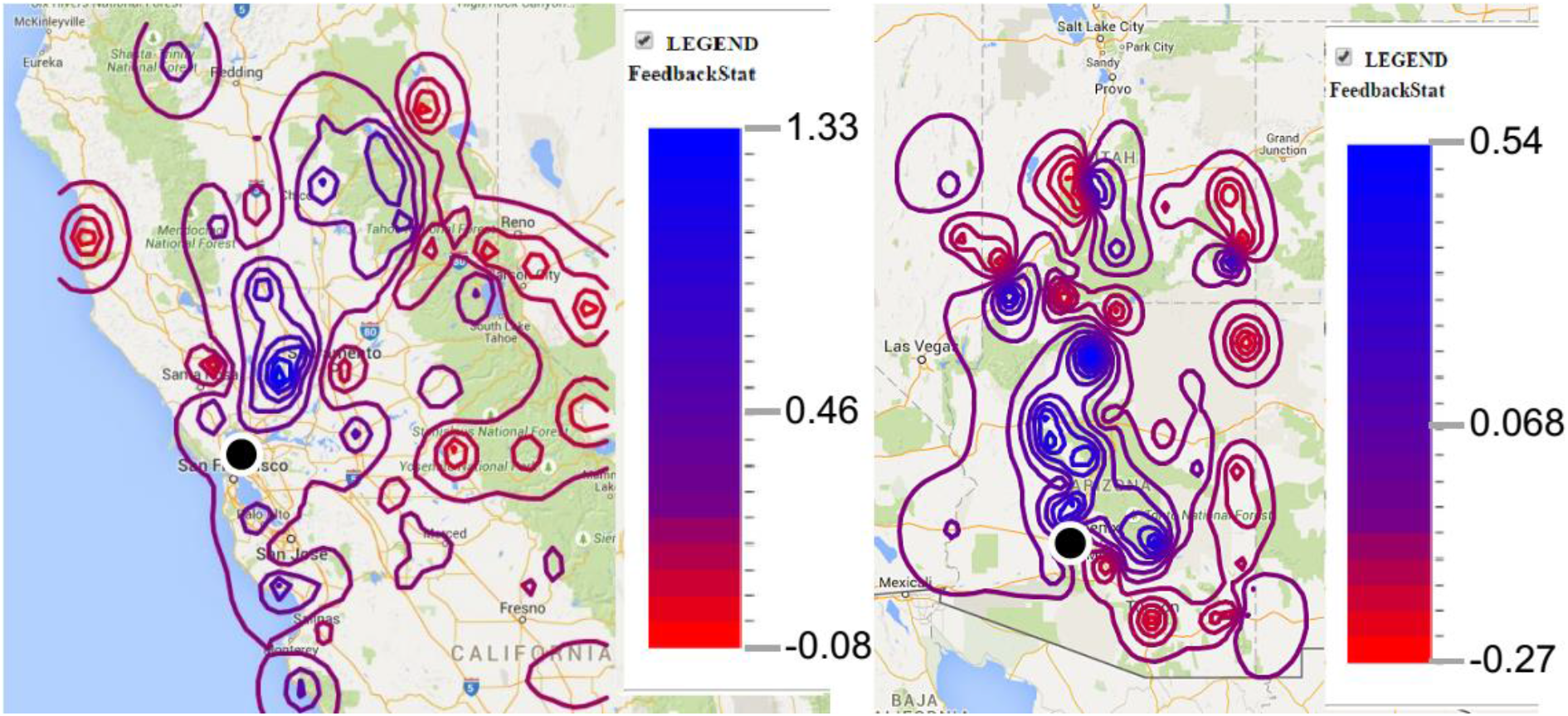
Regions of homogenous ***F*** delimited by kriging in the vicinity of San Francisco (left panel) and Phoenix (right panel) revealing the downwind plumes of positive rainfall feedback. On each panel the city is indicated with a black dot.

One hypothesis could be that urban centers emit some type of aerosol, not described to date, persistently in response to rainfall and that fosters positive feedback. The greater seasonality of rainfall feedback for Phoenix compared to Tucson suggests that differences in land use would give clues to the nature of these aerosols for the Phoenix region if they exist. These observations also raise questions about the dynamics of aerosol composition of urban centers and the synergistic or antagonistic interactions that could occur among the different types of aerosols. This leads to another hypothesis about the effect of urban centers: that urban deactivation of rainfall initiators increases the importance of freshly produced GCCN and INP recurring at intervals following a key day. This hypothesis is compatible with recent observations that aerosols associated with air pollution can enhance precipitation depending on local meteorological conditions, the presence of dust and other factors that lead to particle interactions that favor growth of cloud drops, and location of precipitation (10, 41). Tests of the hypotheses evoked here would require not only the assessment of a range of biological, biogenic and anthropogenic aerosols and dust at selected study sites but also would need careful analyses of the chemical interactions that could alter their cloud-activity

### Hypothesis 4. Pollution from petroleum production fosters negative feedback

As examples of sites with the most negative values of ***F***, sites to the east and south of Houston, Texas are a striking exception to the trend of positive ***F*** near large urban centers. There are major oil refineries at these positions, Port Arthur and Beaumont to the east and Galveston Bay and Deer Park to the south. Fig. 7 shows areas of negative feedback extending to the NW from them. Another large refinery at Baytown is on the eastern edge of the metropolitan area. The Baytown operation has the largest output in the nation and sulfur-rich oils are processed. In 1990, for example, it produced over 50000 tons of SO_2_ per year according to the US Environmental Protection Agency. Although in the immediate vicinity of these refineries there is negative feedback, further downwind, it appears as if the negative effect has been submerged in an area of urban positive feedback that also extends to the NW. This suggests that, as under Hypothesis 3 above, the negative effect of pollution on rainfall was overcome through interaction with urban aerosols in the meteorological context of the Houston region. Interestingly, the outcome of the atmospheric processes in the vicinity of this refinery for rainfall feedback is the same as those occurring in the most northwest corner of Washington where the sources and types of aerosols are probably strikingly different. Both contexts lead to the greatest negative feedback of all sites examined in this work (Fig. 2). By examining the various contexts where there is negative rainfall feedback, a comprehensive list of aerosols, and their concentration and dynamics that are inhibitory to rainfall could be elaborated. In conjunction with tests of hypotheses about aerosol interactions described above, the environmental contexts that alleviate the negative effects could be described.

**Figure 2.**
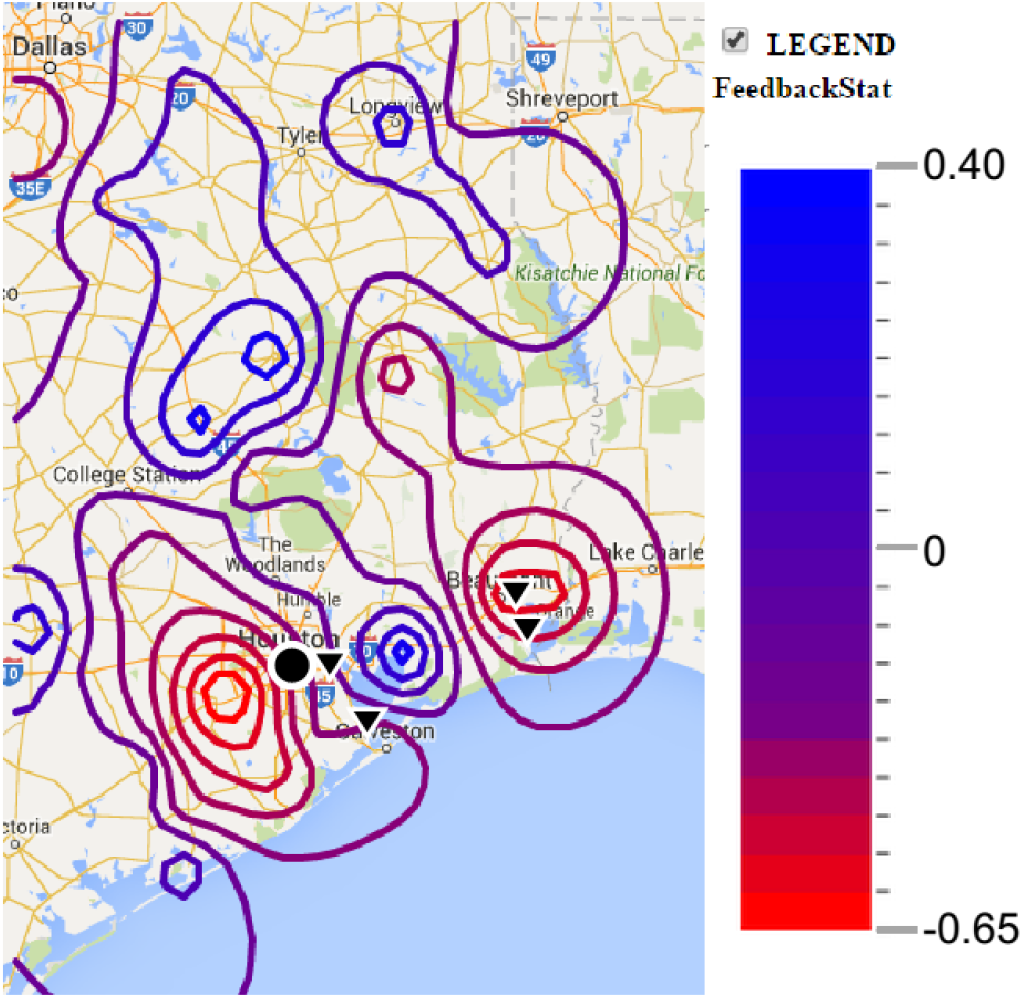
Regions of homogenous ***F*** delimited by kriging in the vicinity of Houston, Texas revealing the plumes of negative rainfall feedback in the vicinity of the petroleum refineries to the east and west of Houston. Refinery complexes are indicated with black triangles and Houston is indicated by the black dot.

## DISCUSSION

The geographic variability of rainfall feedback revealed in rainfall feedback patterns (28) and in the hypotheses we present here illustrates the problem of confounding effects of site-specific traits on the influence that aerosols have on precipitation. Rainfall feedback as defined by our index concerns processes that are apparently amplified in a persistent manner after a rainfall. In some cases the apparent persistence could be due to a series of events, such as fungal sporulation, that start in succession after a rainfall and contribute overall to a persistent increase in certain types of cloud-active aerosols. Persistent increase could also be due to microbial growth that begins near the time of the rainfall. Overall, microorganisms are reasonably the main focus for studies of persistent aerosols because of their rapid response times and growth rates. Exploration of the processes underlying rainfall feedback will foster the discovery of persistent rain-responsive cloud-active aerosols beyond what has been described elsewhere (28). Furthermore, unraveling the processes underlying rainfall feedback will also advance our understanding of the impact of biological aerosols on rainfall and their interaction with non-biological aerosols, in the context of disparate meteorological conditions. In this exploration it should be kept in mind that positive rainfall feedback is not necessarily due to direct enhancement of rainfall by persistent aerosols. It could also be the result of the alleviation of inefficient or inhibitory cloud-activation processes from urban or other pollution sources. Furthermore, it should also be kept in mind that near ground level increases of aerosols such as biological INPs, for example, does not mean that these aerosols are effectively transported into clouds. Means to directly assess microbial flux into the atmosphere or to validate modeled estimations remains a challenging frontier.

Here we present a rationale for developing site-specific hypotheses about the impact of aerosols on meteorological conditions that are ripe for rainfall and about the underlying geographical, seasonal and land-use contexts. This rational can be set into the framework of a set of generic hypotheses depending on whether the trend of rainfall feedback is positive, negative or null. This framework is illustrated in Table 1. We have developed this framework and the associated tools to contribute to the design of field experiments finely tuned to site-specific hypotheses and thereby to a more refined understanding of the context and the extent to which aerosols influence the fate of cloud processes.

**Table 1.** Generic hypotheses and questions relative to rainfall feedback trends.

| <u>Rainfall feedback trend<sup>1</sup>:</u> | <u>Hypotheses (H) and Questions (Q)</u> |
| --- | --- |
| Positive | **H<sub>1</sub>:** Sources of rain-responsive aerosols are present but the quantity of these aerosols is a limiting factor for rainfall prior to a rain event.<br>**H<sub>2</sub>:** Cloud-active aerosols are not limiting in abundance before a rain event, but they are inefficient and their activity is enhanced after a rain event.<br>**Q:** Are there conditions under which the quantity and activity of these aerosols could increase to levels that are inhibitory to precipitation? |
| Negative | **H<sub>1</sub>:** Cloud-active aerosols are abundant prior to a rain event. Sources of rain-responsive aerosols are present and the increase in their abundance after rainfall leads to a total atmospheric load of cloud-active aerosols that is inhibitory to precipitation.<br>**H<sub>2</sub>:** Cloud-active aerosols are abundant prior to a rain event but of low efficiency. Rainfall sets off processes that increase the efficiency of these cloud-active aerosols leading to a total atmospheric load of cloud-active aerosols that is inhibitory to precipitation.<br>**Q<sub>1</sub>:** To what extent can aerosol abundance be inhibitory to rainfall?<br>**Q<sub>2</sub>:** Would elimination of processes that inhibit the activity of rain-responsive aerosols lead to positive rainfall feedback? |
| Null | **H<sub>1</sub>:** Sources of rain-responsive aerosols are absent.<br>**H<sub>2</sub>:** Rain-responsive aerosols are inactivated after their formation.<br>**Q:** Would rain-responsive aerosols have effects on precipitation if they were introduced or if the inactivation processes were inhibited? |

<sup>1</sup>These refer to regional trends that are stable or that alter with time and/or season of the year.

## Acknowledgements

This work was supported by in-house funds from INRA and Montana State University and the personal resources of E. K. Bigg.

## Footnotes

1 For online information about the circulation of air masses in Arizona : http://www.library.arizona.edu/exhibits/swetc/azso/body.1_div.3.html, http://ag.arizona.edu/extension/riparian/pub/UARA_07-17-07_chapter6.pdf.

## References

1. AghaKouchak A, Feldman D, Hoerling M, Huxman T, Lund J. 2015. Water and climate: Recognize anthropogenic drought. Nature 524: 410–1

2. Angassa A, Oba G. 2013. Cattle herd vulnerability to rainfall variability: responses to two management scenarios in southern Ethiopia. Tropical Animal Health and Production 45: 715–21

3. Bigg EK. 1958. A long period fluctuation in freezing nucleus concentrations. J. Meteorology 15: 561–2

4. Bigg EK. 2004. Gas emissions from soil and leaf litter as a source of new particle formation. Atmospheric Research 70: 33–42

5. Bigg EK, Soubeyrand S, Morris CE. 2015. Persistent after-effects of heavy rain on concentrations of ice nuclei and rainfall suggest a biological cause. Atmos. Chem. Phys. 15: 2313–26

6. Creamean JM, Ault AP, Ten Hoeve JE, Jacobson MZ, Roberts GC, Prather KA. 2011. Measurements of aerosol chemistry during new particle formation events at a remote rural mountain site. Environmental Science & Technology 45: 8208–16

7. Dettinger MD, Ralph FM, Das T, Neiman PJ, Cayan DR. 2011. Atmospheric rivers, floods and the water resources of California. Water-Sui 3: 445–78

8. Devlin C, Hendrix CS. 2014. Trends and triggers redux: Climate change, rainfall, and interstate conflict. Political Geography 43: 27–39

9. Eversmeyer MG, Kramer CL. 2000. Epidemiology of wheat leaf and stem rust in the Central Great Plains of the USA. Annual Review of Phytopathology 38: 491–513

10. Fan J, Leung LR, DeMott PJ, Comstock JM, Singh B, et al. 2014. Aerosol impacts on California winter clouds and precipitation during CalWater 2011: local pollution versus long-range transported dust. Atmos. Chem. Phys. 14: 81–101

11. Fjelde H, von Uexkull N. 2014. Climate triggers: Rainfall anomalies, vulnerability and communal conflict in Sub-Saharan Africa. Political Geography 31: 444–53

12. Friesen TL, de Wolf ED, Frankl LJ. 2001. Source strength of wheat pathogens during combine harvest. Aerobiologia 17: 293–9

13. Geils BW, Hummer KE, Hunt RS. 2010. White pines, Ribes, and blister rust: a review and synthesis. Forest Pathology 40: 147–85

14. Givati A, Rosenfeld D. 2004. Quantifying precipitation suppression due to air pollution. J. Appl. Meteorol. 43: 1038–56

15. Grossmann EB, Kagan JS, Ohmann JA, May H, Gregory MJ, Tobalske C. 2008. The Pacific Northwest regional GAP analysis project: Final report on Land Cover Mapping Methods, Map Zones 2 and 7, PNW ReGAP. Institute for Natural Resources, Oregon State University,Corvalis, OR, USA. 66 pp.

16. Hallett J, Mossop SC. 1974. Production of secondary ice particles during the riming process. Nature 249: 26–8

17. Hamilton LM, Stakman EC. 1967. Time of stem rust appearance on wheat in the western Mississippi basin in relation to the development of epidemics from 1921 to 1962. Phytopathology 57: 609–14

18. Hobbs PV, Radke LF, S.E. S. 1970. Cloud condensation nuclei from industrial sources and their apparent influence on precipitation in Washington state. J. Atmos. Sci. 27: 81–9

19. Huffman JA, Prenni AJ, DeMott PJ, Pöhlker C, Mason RH, et al. 2013. High concentrations of biological aerosol particles and ice nuclei during and after rain. Atmospheric Chemistry and Physics 13: 6151–64

20. Letcher T, Cotton WR. 2014. The effect of pollution aerosol on wintertime orographic precipitation in the Colorado Rockies using a simplified emissions scheme to predict CCN concentrations. Journal of Applied Meteorology and Climatology 53: 859–72

21. Levin Z, Cotton WR, eds. 2008. Aerosol Pollution Impact on Precipitation: A Scientific Review: Springer Netherlands. 386 pp.

22. Lindemann J, Upper CD. 1985. Aerial dispersal of epiphytic bacteria over bean plants. Appl. Environ. Microbiol. 50: 1229–32.

23. Lindow SE, Arny DC, Upper CD. 1978. Distribution of ice nucleation-active bacteria on plants in nature. Appl. Environ. Microbiol. 36: 831–8

24. Luk KC, Ball JE, Sharma A. 2000. A study of optimal model lag and spatial inputs to artificial neural network for rainfall forecasting. Journal of Hydrology 227: 56–65

25. Merikanto J, Spracklen DV, Mann GW, Pickering SJ, Carslaw KS. 2009. Impact of nucleation on global CCN. Atmos. Chem. Phys. 9: 8601–16

26. Morris CE, Leyronas C, Nicot PC. 2014. Movement of Bioaerosols in the Atmosphere and the Consequences for Climate and Microbial Evolution (Chapter 16). In Aerosol Science: Technology and Applications, ed. I Colbeck, M Lazaridis, pp. 393–416: John Wiley & Sons, Ltd.

27. Morris CE, Sands DC, Glaux C, Samsatly J, Asaad S, et al. 2013. Urediospores of rust fungi are ice nucleation active at > −10 °C and harbor ice nucleation active bacteria. Atmos. Chem. Phys. 13: 4223–33

28. Morris CE, Soubeyrand S, Bigg EK, Creamean JM, Sands DC. 2016. Mapping rainfall feedback to reveal the potential sensitivity of precipitation to biological aerosols. Bull. Amer. Meteor. Soc. (***under revision***)

29. Neiman PJ, Ralph FM, Moore BJ, Hughes M, Mahoney KM, et al. 2013. The landfall and inland penetration of a flood-producing atmospheric river in Arizona. Part I: Observed synoptic-scale, orographic, and hydrometeorological characteristics. Journal of Hydrometeorology 14: 460–84

30. Neiman PJ, Ralph FM, Wick GA, Lundquist JD, Dettinger MD. 2008. Meteorological characteristics and overland precipitation impacts of atmospheric rivers affecting the west coast of North America based on eight years of SSM/I satellite observations. Journal of Hydrometeorology 9: 22–47

31. Pfender W, Graw R, Bradley W, Carney M, Maxwell L. 2006. Use of a complex air pollution model to estimate dispersal and deposition of grass stem rust urediniospores at landscape scale. Agric. For. Meteorol. 139: 138–53, doi:10.1016/j.agrformet.2006.06.007

32. Pielke RA, Adegoke J, Beltran-Przekurat A, Hiemstra CA, Lin J, et al. 2007. An overview of regional land-use and land-cover impacts on rainfall. Tellus B 59: 587–601

33. Redford AJ, Bowers RM, Knight R, Linhart Y, Fierer N. 2010. The ecology of the phyllosphere: geographic and phylogenetic variability in the distribution of bacteria on tree leaves. Environmental Microbiology 12: 2885–93

34. Riccobono F, Schobesberger S, Scott CE, Dommen J, Ortega IK, et al. 2014. Oxidation products of biogenic emissions contribute to nucleation of atmospheric particles. Science 344: 717–21

35. Rosenfeld D. 2000. Suppression of rain and snow by urban and industrial air pollution. Science 287: 1793–6

36. Rosenfeld D, Woodley WL, Axisa D, Freud E, Hudson JG, Givati ACD. 2008. Aircraft measurements of the impacts of pollution aerosols on clouds and precipitation over the Sierra Nevada. Journal of Geophysical Research: Atmospheres 113: doi:10.1029/2007JD009544

37. Stopelli E, Conen F, Morris CE, Hermann E, Bukowiecki N, Alewell C. 2015. Ice nucleation active particles are efficiently removed by precipitating clouds. Scientific Reports 5:16433, DOI:10.1038/srep16433

38. Tan L, Cai Y, An Z, Cheng H, Shen C-C, et al. 2015. A Chinese cave links climate change, social impacts, and human adaptation over the last 500 years. Scientific Reports 5: 12284

39. Toth E, Brath A, Montanari A. 2000. Comparison of short-term rainfall prediction models for real-time flood forecasting. Journal of Hydrology 239: 132–47

40. Willems P, Arnbjerg-Nielsen K, Olsson J, Nguyen VTV. 2012. Climate change impact assessment on urban rainfall extremes and urban drainage: Methods and shortcomings. Atmospheric Research 103: 106–18

41. Xiao H, Yin Y, Jin L, Chen Q, Chen J. 2015. Simulation of aerosol effects on orographic clouds and precipitation using WRF model with a detailed bin microphysics scheme. Atmospheric Science Letters 15: 134–9

42. Zeppel MJB, Wilks JV, Lewis JD. 2014. Impacts of extreme precipitation and seasonal changes in precipitation on plants. Biogeosciences 11: 3083–93

